# A ChatGPT Assisted Reading Protocol for Undergraduate Research Students

**DOI:** 10.1101/2024.09.11.612473

**Authors:** Marcus Sambar, Gonzalo R. Vázquez, Anne V. Vázquez, Frank X. Vázquez

**Affiliations:** Department of Chemistry, St. John’s University, Queens, NY 11439, USA

**Keywords:** undergraduate research, reading, scientific literature, AI, ChatGPT

## Abstract

A major challenge for undergraduate students is reading scientific literature. This is especially true in biophysics, where many of the concepts may not have been covered in undergraduate courses. Students can become overwhelmed, which may lead to less overall engagement with the scientific literature. In response, we have developed a guided reading protocol that combines pre-reading strategies, structured notetaking, and ChatGPT to help students clarify unfamiliar concepts in an interactive way. To test the protocol, participants in this study were given an initial survey to determine their experience with reading scientific literature. After this they were given an abridged biophysics paper and the protocol. The ChatGPT transcripts were analyzed using open coding and the students were given a post-study survey. We found most students did not appear to regularly engage with the literature, possibly due to content barriers they encountered. Analyzing their transcripts, we observed that students asked for definitions, explanations, summaries, and simplifications. Overall, students reported that using ChatGPT was a positive experience and they expected to use ChatGPT in the future. From this work, we expect this new protocol may be a way to keep novice students from becoming discouraged when reading scientific papers and keep them engaged with the current literature.

## Introduction

When a student joins a research lab they encounter scientific literature, often for the first time, that goes beyond what they have encountered in their coursework. Undergraduates engaged in biophysical research, especially, must grapple with structural biology and biophysics papers with unfamiliar terms, concepts, and methodologies. Encountering these new ideas as well as the complexity and unique format of scientific literature act as barriers, preventing students from engaging with the literature in a meaningful way. Without supports and guidance, reading literature may cause anxiety that can hinder the learning experience for the undergraduate researcher.^1^ Gaining competency in reading literature is necessary not only for developing scientific skills but also contributes to improving student motivation and sense of belonging in their field, which is critical for retaining students in the sciences.^2-6^

Although this is an important topic for training students to be scientists, limited studies on how undergraduate STEM students approach reading scientific literature have been reported. A study by Lennox and colleagues indicated that undergraduates use multiple approaches to finding evidence in reading primary scientific literature, such as prioritizing different sections of the paper, using figures and text to support conclusions, and skimming or even skipping numerical data presented in the text.^7^ While most students were able to identify the main points of a given text, they struggled to provide evidence for their conclusions. A study by Mitra and colleagues observed that many undergraduate STEM students struggled to identify the research hypothesis of articles while participating in a guided literature reading curriculum.^8^ Additionally, studies observed a disconnect between students’ self-perception of their scientific reading comprehension ability and their actual skills,^7, 8^ which may contribute to undergraduate students not fully engaging with literature because they do not realize that they are not doing so effectively. Reading scientific literature requires different skills than “everyday” reading,^6^ such as being able to prioritize which section to read, understanding the context of quantitative data or figures with relation to the text, and knowing how to decipher technical terms and jargon. As a result, scientific paper skills need to be explicitly taught to undergraduate research students along with the use of new technologies, such as ChatGPT, to help students fully engage with the material.

### Scientific and Pedagogic Background

Interventions have been developed and assessed to help undergraduate STEM students engage with scientific literature more effectively. In a project by Round and colleagues, a template called “figure facts” was used to help students read and interpret scientific literature by having students focus on experimental data presented in an article’s figures. Students showed improvement in data interpretation and attitudes about reading scientific literature.^2^ Mitra and colleagues developed a curriculum to guide undergraduates’ reading of scientific articles by focusing on the hypothesis and conclusions drawn from experimental results using a series of guided worksheets and workshops. They found that students’ confidence improved with the curriculum.^8^ Gold and colleagues embedded a scientific literature reading and graph interpretation curriculum into a ten week REU program. They analyzed the reading strategies used via survey and eye-tracking studies to compare REU participants to experts, and found that with the embedded curriculum and authentic research experience in the REU, participants made gains in their reading strategies and graph interpretation skills.^6^ To advance undergraduates’ scientific literacy and understanding of the publication process, Otto and colleagues developed a curriculum in which students peer reviewed preprints, which was found to improve students’ scientific literacy and sense of belonging in STEM.^9^

These interventions have greatly helped students’ understanding of the format and reading of scientific literature. However, conventional methods are unable to address the issue of encountering unfamiliar terms within the context of a research article in a timely manner, so as to not frustrate and discourage the student. This is especially important when undergraduates are doing research and reading journal articles in a multidisciplinary field like biophysics, where they need to engage with terms that are outside the scope of the material they have learned in their classes. This study aims to show that a guided reading protocol that includes artificial intelligence (AI) based Large Language Models (LLMs) such as ChatGPT can be used to bridge knowledge gaps and help undergraduate research students effectively read and understand scientific literature.

The advent of LLMs like ChatGPT has quickly changed how individuals interact with information. These new tools can be used for answering questions, summarizing information, and can play a role in improving equity by minimizing language barriers for diverse learners.^10-13^ Many interesting use cases for AI in postsecondary science education are already being explored and analyzed in the literature. A common theme in published studies is the importance of training students in AI literacy so that they understand how to use a tool such as ChatGPT effectively and understand its limits.^4, 10, 14-18^ Further, much of the literature has highlighted the need to develop training for students in prompt engineering, the process of crafting AI queries to elicit better answers.^13-17^ Tassoti developed a structured framework for posing questions to ChatGPT in a way that led to satisfactory responses without needing to copy large sections of text into the chat.^16^ While nearly all studies stress the importance of prompt engineering, studies on the use of LLMs in STEM education vary in how the students are interacting with the chatbot. In one study by Exintaris and colleagues, students were introduced to problem-solving methods, taught a metacognitive scaffolding approach for problem solving, and then asked to critique a ChatGPT generated solution to a problem. Most students were able to find flaws in the LLM-generated results, showing that with scaffolding students were able to think critically about the responses provided by ChatGPT.^19^ A study by Pence and colleagues involved analyzing a three-part assignment for undergraduate chemistry students: students used ChatGPT to summarize an article from a chemistry trade magazine, analyzed ChatGPT output for a numerical solution, and used ChatGPT to analyze primary scientific literature by using it to answer specific questions. In summarizing the trade magazine article, the students largely felt this was a good use of the LLM, and that the summaries were generally accurate but superficial. For the assignment involving using ChatGPT to answer questions from primary scientific literature, results were mixed. Students became frustrated with prompting, and some hallucinations and superficial answers occurred. However, when analyzed by faculty it was found that the majority of responses from ChatGPT were at least partially correct.^20^

To facilitate effective engagement with scientific literature for undergraduate research students, we have developed a new approach that combines pre-reading strategies adapted from the book, *Teaching Students to Learn* by McGuire^21^, notetaking, and the use of artificial intelligence-based Large Language Models (LLMs) like ChatGPT. The LLM assists students in clarifying unfamiliar terms and concepts, preventing these from becoming barriers, thus enabling students to better integrate the paper’s content into their own research.

## Methods

Research subjects were recruited via an email from the pool of undergraduate students participating in research in the chemistry department at a primarily undergraduate institution. The students ranged from second to fourth year students with less than one year to two years of undergraduate research experience, with majors in chemistry, biochemistry, and biology. A pre-intervention survey was given to assess the students’ prior experience and comfort level with reading scientific literature and with using AI chatbots such as ChatGPT.

After completing the initial survey, students were sent two items: a ChatGPT-assisted literature reading protocol and the title, authors, abstract, and introduction of the 2018 paper “Stabilized Coronavirus Spikes are Resistant to Conformational Changes Induced by Receptor Recognition or Proteolysis” by Kirchdoerfer *et al* ^22^. This structural biology paper was chosen because we expected it to present unknown terms and methodologies to chemistry or biology research students, but the coronavirus subject matter would still provide some familiarity. The paper was abridged to these sections because the introduction is often read first by students and it is where many unfamiliar terms and concepts that may act as barriers would be presented.

The ChatGPT-assisted literature reading protocol consisted of four elements to guide the students in using the chatbot as a tool for understanding scientific literature. This procedure is also included in the supplemental information as Figure S1.

1. **<u>Previewing the paper</u>**. The reader is encouraged to analyze the title, abstract, headings, and figures to gain meaning.
  a. <u>Analyzing the title</u>. The reader is asked to pick out key words that give the paper’s main focus.
  b. <u>Analyzing the abstract</u>. The reader is encouraged to analyze the abstract for (1) key findings and (2) the general format of an abstract: importance, problem to be addressed, approach, and insights.
  c. <u>Analyzing headings and figures</u>. The reader is also asked to skim text headings and figures to anticipate the questions the paper will answer. It should be noted that the abridged paper we provided did not in this case include a figure.
2. **<u>Guided reading of the paper</u>**. In reading the paper, the reader is told to highlight key text with an erasable method and to refine/edit highlighting as they read. The reader is also asked to make margin notes of important points and if needed, extended notes in a separate notebook.
3. **<u>Using ChatGPT for clarifications</u>**. The reader is told to use ChatGPT to help clarify unfamiliar concepts, terms, or methodologies that feel like stumbling blocks, and are walked through how to effectively use ChatGPT for this purpose:
  a. <u>Introduce yourself</u>. First, the reader introduces themselves with their academic and research background and any other pertinent information.
  b. <u>Introduce the paper</u>. The reader informs ChatGPT about the paper, giving the paper’s title and general field.
  c. <u>Ask questions</u>. The reader asks questions to ChatGPT and is encouraged to ask clarifying or follow-up questions as needed.
4. **<u>Post-reading summary</u>:** Lastly, the reader is encouraged to review their notes and ChatGPT transcript to write a summary of the paper. The summary can be as brief as one or two sentences or bullet points to synthesize ideas and serve as a record of the paper for the reader.

This protocol was designed to be a general procedure for reading papers, and as a result it does include instructions in the previewing step directing students to analyze headings and figures. Because this was an initial pilot study, we chose to focus on the introduction, which did not include figures. Our goal was to explore generally how students interacted with the chatbot in this first study. In future work, we will be testing the protocol on an entire paper, which will give us insight into how students interact with figures in the paper.

Students were asked to save their ChatGPT transcripts as PDF files and send them to researchers via email upon completion of the guided reading task. The ChatGPT transcripts were anonymized prior to analysis. Students were then sent a post-implementation survey about their experience using the ChatGPT-assisted literature reading protocol. We did not ask students which version of ChatGPT they used; however, we do not believe this had an impact on this study. The fundamental functions needed for this protocol – interactivity, summarizing, and explaining – are available across all versions of the chatbot.

Responses to multiple selection and Likert-scale questions on the pre-and post-implementation surveys were tallied, and free response questions were coded using open coding for common themes.^23^ ChatGPT transcripts were analyzed in two ways. First, the number of questions asked by each student was tallied. Second, open coding was used to analyze the types of questions being asked to ChatGPT. Five question-type codes emerged. Questions corresponding to these codes were tallied.

## Results and Discussion

### Pre-implementation survey

In the pre-implementation survey, students were asked about their experience with and perceptions about scientific literature. The survey questions that students were asked, along with their responses to the survey are included in Supplemental File 1. Fourteen students responded to this survey. The first part of the survey asked about experience reading journal articles, and the second part of the survey asked about approaches to reading journal articles and previous experience using LLMs such as ChatGPT.

The first set of questions were focused on understanding students experience and comfort with reading scientific papers (Figure 1). These students were asked whether their research advisors or classes taught them how to read papers, how confident they felt about reading papers, and how often they actually read papers. Students reported mixed views on how well they felt their research mentors and course instructors prepared them to read scientific literature, with 43% (6 of 14) of students agreeing or strongly agreeing that research faculty had taught them to read scientific literature, while 50% (7 of 14) felt the same about their courses. Students were also asked to rank their confidence in reading scientific literature, with 57% (8 of 14) of respondents saying they felt confident or very confident and 29% (4 of 14) saying they felt moderately confident reading literature. However, this self-reported measure did not correspond with students’ actual limited experience reading literature, in agreement with previous studies showing that undergraduate students often overestimate their reading skills.^7, 8^ Only 43% (6 of 14) of students agreed or strongly agreed that they read papers regularly. When asked how many journal articles students read per month related to their research project, 29% (4 of 14) of students reported not reading any journal articles, 64% (9 of 14) reported reading 1-2 articles, and only 7% (1 of 14) reported reading 3-4 articles (Figure 2). When asked how many articles they read per month in their classes, 57% (8 of 14) students reported not reading any, 21% (3 of 14) reported reading 1-2 articles, 7% (1 of 14) reported reading 3-4 articles, and 14% (2 of 14) reported reading more than 4 articles.

**Figure 1:**
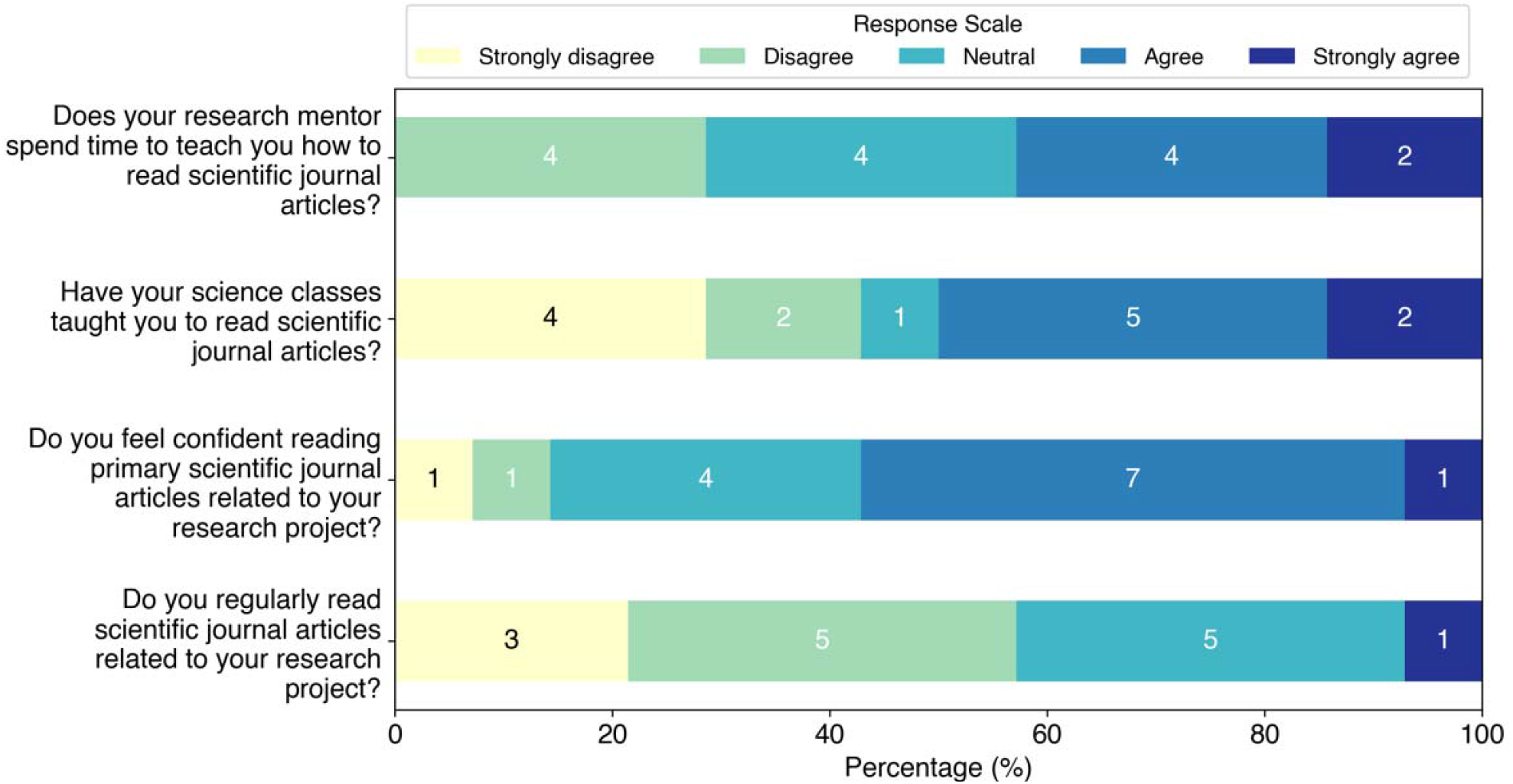
Responses to questions about student experience and confidence reading scientific literature.

**Figure 2:**
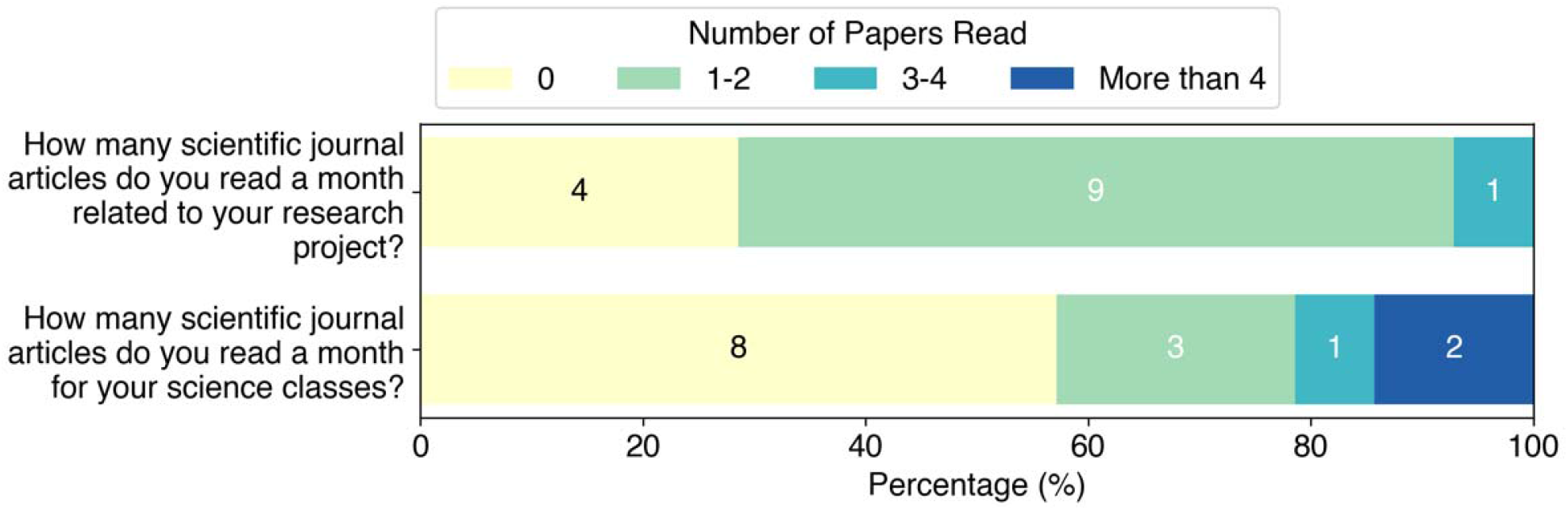
Student responses to how many papers they read per month for research or science classes.

When asked about approaches students take to reading scientific literature, allowing for multiple responses (Figure 3), the most common approach was reading the abstract first (64%, 9 of 14) followed by reading the article from beginning to end (50%, 7 of 14), and jumping around the article (50%, 7 of 14). Interestingly, no students responded that they read the conclusions first. Students were also asked to provide free responses to describe reading strategies they have been taught. The most common strategy students reported was to read the abstract first and then look at the figures (36%, 4 of 11). We also asked students what they do when they encounter an unfamiliar word, phrase, or concept, allowing for multiple responses (Figure 4). The most common response was using a search engine such as Google (93%, 13 of 14), followed by using LLMs such as ChatGPT (50%, 7 of 14). We can see that undergraduate student researchers are using online resources to decode unfamiliar terms and concepts, but generally without specific guidance that would help them do this effectively and efficiently.

**Figure 3:**
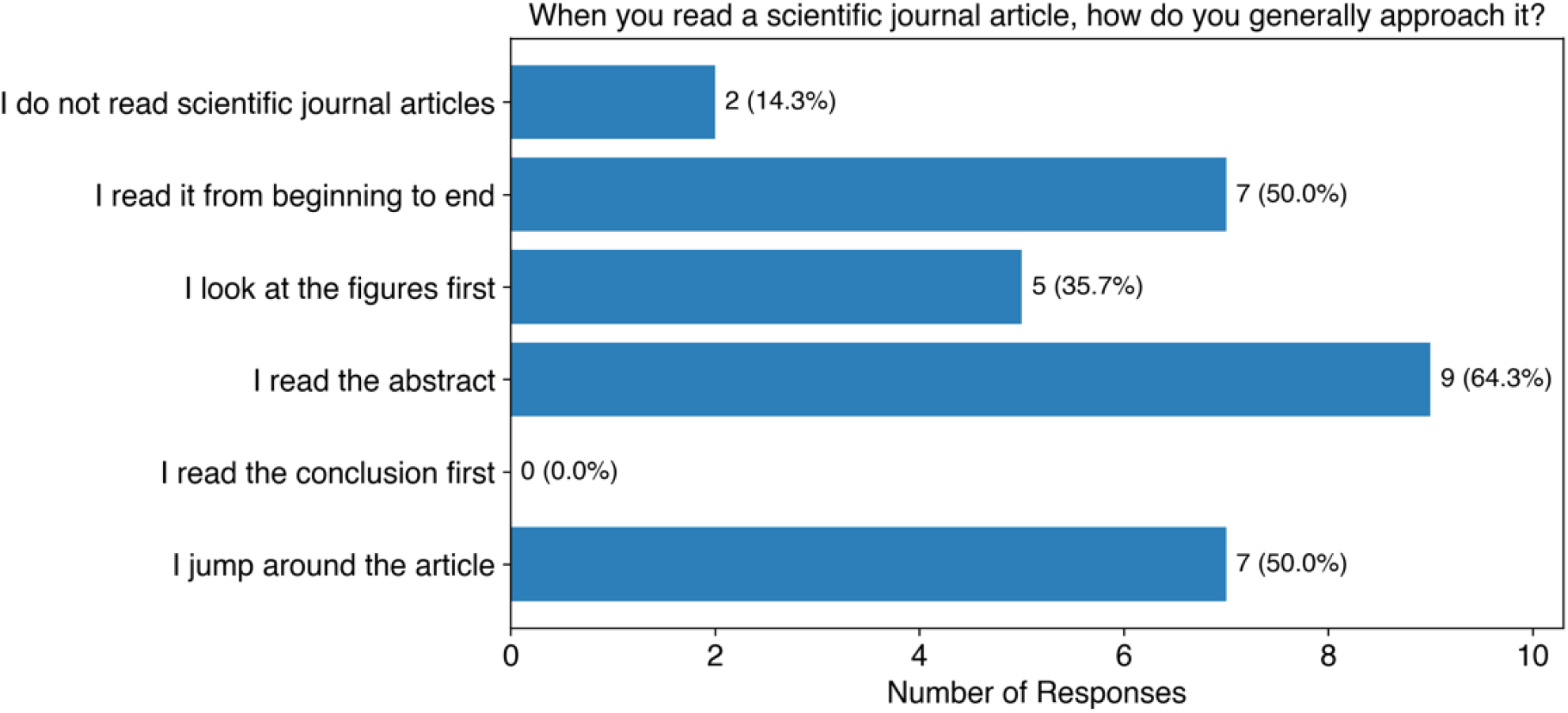
Student survey responses on how they approach reading scientific journal articles. Students were allowed to pick multiple responses.

**Figure 4:**
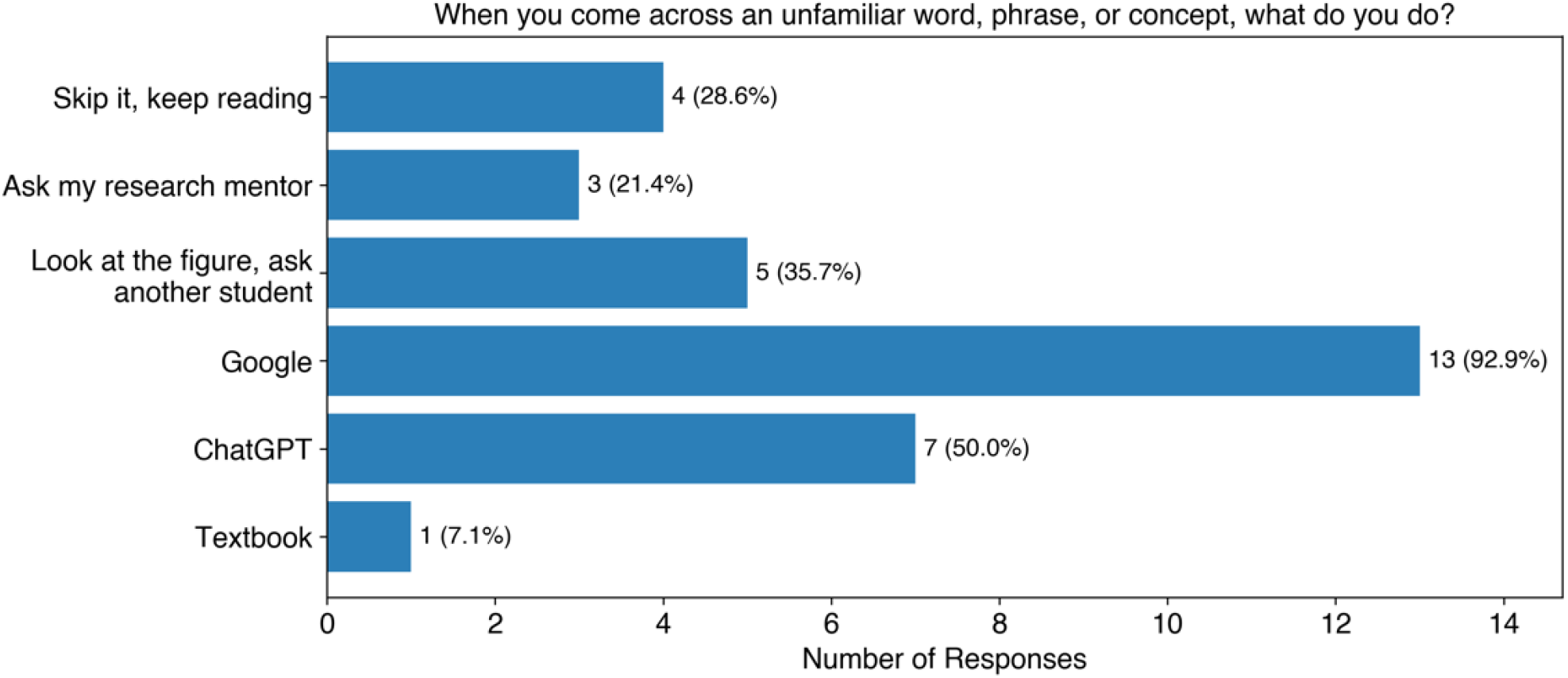
Student responses to how they respond when encountering unknown concepts in scientific journal articles. The approaches have been abridged for figure clarity. The full survey question can be found in Supplemental File 1.

In a write-in response question, the students were asked what they perceive to be the most difficult aspect of reading scientific literature. Two common themes emerged: encountering unfamiliar vocabulary, terminology, and abbreviations (46%, 6 of 13) and deciphering scientific writing in terms of its length, density, and style (54%, 7 of 13). While the undergraduate researchers in this study expressed confidence in their ability to read scientific literature, they also indicated a lack of experience and instruction on how to approach reading journal articles.

Undergraduate students appear to be employing tools such as search engines and LLMs on their own to tackle the challenges of reading scientific literature, but still perceive significant barriers to their understanding. This suggests that training research students to use LLMs such as ChatGPT can be a productive tool for removing the obstacles of unfamiliar terminology. It may also be beneficial for undergraduates who are tackling difficult scientific literature, possibly for the first time.

### ChatGPT Transcripts

A total of eleven students completed the ChatGPT-assisted reading of the article and returned the ChatGPT transcripts for analysis. Anonymized ChatGPT transcripts were analyzed for (1) number of interactions initiated by each student and (2) common types of questions and statements that were posed by the students. Here, we counted interactions as either giving background information to ChatGPT or asking a specific question. The average number of interactions with ChatGPT was 7.1 interactions, with a range of 5 to 14 interactions in each transcript.

To investigate how undergraduate research students were interacting with ChatGPT, open coding was used to group the types of questions and statements posed into five common themes, a description of the student’s educational background, definitions of unfamiliar terms, requests for an explanation, asking the chatbot to summarize text or the conversation, and simplifications of the material The number of instances of each of these themes is summarized in Figure 5 and further discussed in the text below.

**Figure 5:**
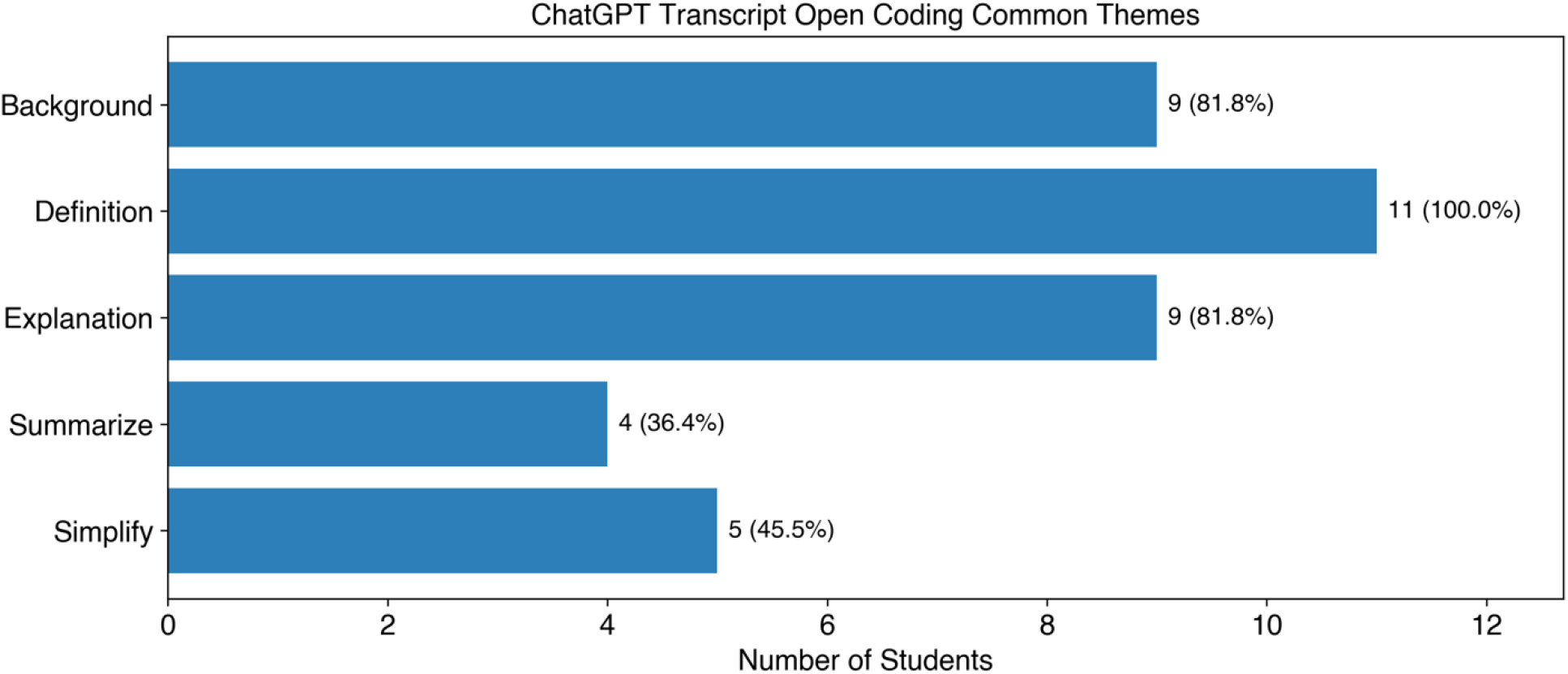
Number of students using the common interactions determined from open coding of student ChatGPT transcripts.

**Figure 6:**
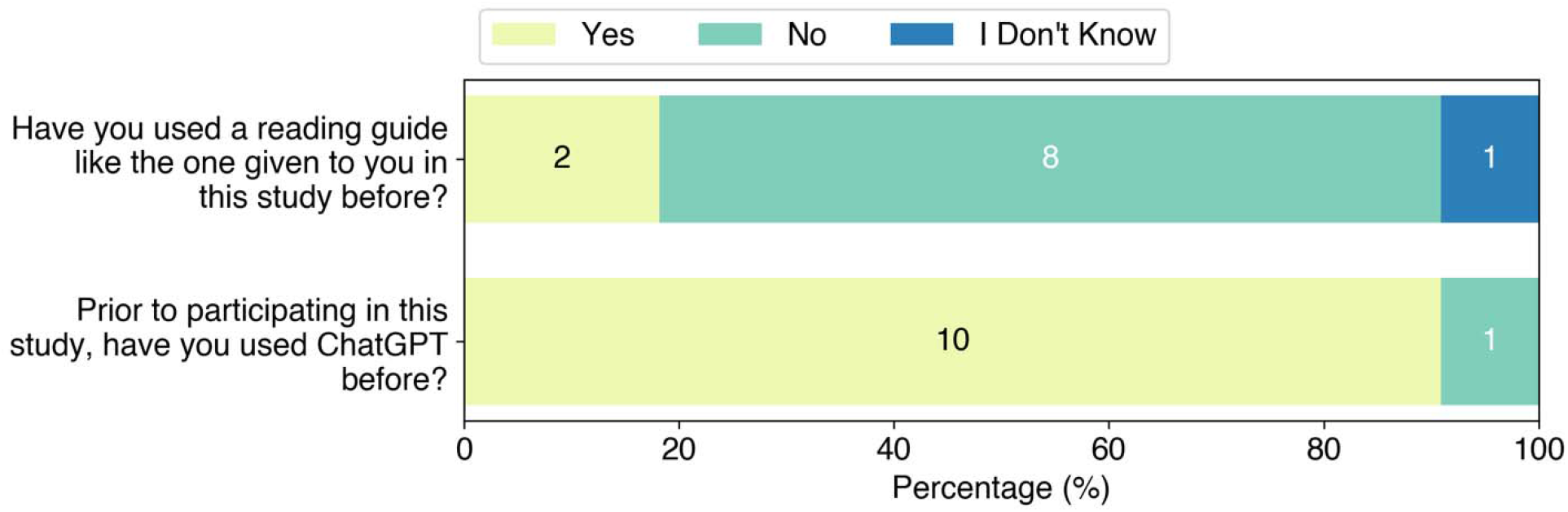
Survey of students’ previous experience with reading protocols and ChatGPT.

1. **Background** The student gave information about their educational background as they were prompted to do in the user guide (82%, 9 of 11 students used this type of interaction). In giving background information, students gave information about their educational background, and some gave information about their experience reading scientific journal articles. Students were prompted to provide information about their academic background in the provided reading guide, to allow ChatGPT to tailor its response to the individual’s educational level. All students who gave background information described their educational background in terms of major and relevant courses completed. For example: *“I’m currently a third-year chemistry student in my second semester. I’ve taken two semesters of general chemistry and organic chemistry, each with analytical chemistry and quantum mechanics classes. I am enrolled in a thermodynamics class right now. I have a very light background in biology, mostly pertaining to protein structures, but chemistry is more of my strong suit*.*”* Of the students who gave background information, 44% (4 of 9) also gave information about their research interests and experience, which was in addition to what they were prompted to do in the reading guide. For example: *“I am a third-year college student majoring in biology. I have taken courses such as general chemistry, organic chemistry, and biology courses such as microbiology and now biochemistry. I also currently do research in a chemistry lab doing analytical chemistry comprised of measuring absorbance using a UV-Vis spectrometer. I rarely read scientific papers and it takes me a long time to thoroughly comprehend even a few sentences*.*”*
2. **Definition** The student asked for the definition of an unfamiliar term (100%, 11 of 11 students used this type of interaction). All students asked definition questions pertaining to unfamiliar words and phrases in the text. We saw in the pre-implementation survey that unfamiliar terms are common stumbling blocks in reading scientific literature. Using an LLM to obtain definitions quickly without having to evaluate multiple sources may help undergraduate researchers to persevere in reading scientific literature quickly and help keep them from feeling discouraged when encountering new concepts. Example definition questions included: *“What does the word “zoonotic” mean?”* *“What is single-particle cryo-EM and what does an ectodomain do?”*
3. **Explanation** The student asked for an explanation about a concept from the text (82%, 9 of 11 students used this type of interaction). In these questions, students asked for an explanation rather than just a definition about the text itself. Explanation questions frequently came after a definition question as the subject aimed to further understand a new term or idea within the context of the definition they received. Examples include: *“Why is it important to prevent the transition from pre-fusion to post-fusion states for the stabilized coronavirus spikes?”* *“Why are coronavirus spike proteins trimeric? Does this contribute something important to their function?”*
4. **Summarize** The student asked for summary of information from the text or summary of ChatGPT response (36%, 4 of 11 students used this type of interaction). Summarize-type questions posed by students took two different forms. Most students gave either the digital object identifier (DOI) of the paper or large quotes from the paper and asked for a summary of the text. This may have been done to lower cognitive load for the reader when encountering scientific literature on an unfamiliar subject. An example of this is: *“The paper I need help understanding is ‘Stabilized coronavirus spikes are resistant to conformational changes induced by receptor recognition or proteolysis’ (DOI: 10*.*1038/s41598-018-34171-7). Could you please summarize the abstract?”* Students also used summarize-type questions to give a summary of their understanding and prompt ChatGPT to respond and add clarification to the summary in order to critique and deepen their own understanding. An example is: *“I have just completed reading the abstract and introduction of Stabilized coronavirus spikes are resistant to conformational changes induced by receptor recognition or proteolysis, a biochemistry journal article. The intro gives the background behind the experiment and its importance while the abstract summarizes the article in a brief paragraph. The intro discusses coronavirus strands like MERS-CoV and SARS-CoV but focuses on the latter. The virus contains large trimeric S-glycoproteins, which aid in recognizing host receptors and virla* [sic] *cell entry. As SARS-CoV breaks down into subunits S1 and S2, with S2 having the S2 cleavage site and S1 with N terminal domains and receptor binding domain. Using cryo-EM analysis, the researchers were able to develop to stop* [sic] *the transition of pre to post fusion states*.*”* From this example, after the student shared their summary, ChatGPT provided definitions of many of the terms the student used (such as subunits S1 and S2) without prompting, and the student replied with follow-up definition and explanation questions. This interaction allowed the student to check and organize their understanding of the material and receive details and feedback from ChatGPT.
5. **Simplify** The student asked ChatGPT to either simplify language from the journal article or simplify its own explanation in terms of language used or length of explanation (46%, 5 of 11 students used this type of interaction). Students approached asking for simplifications in different ways. Some primed ChatGPT in their introductions to use simple language in its responses. These students may have recognized that complex language and unfamiliar terms are a barrier for their own understanding of scientific literature before reading the article. For example: *“I am going to be using your help to analyze and understand a journal article about coronavirus. The article is titled ‘Stabilized coronavirus spikes are resistant to conformational changes induced by receptor recognition or proteolysis*.*’ When you explain things please keep it simple and understandable*.*”* Other students requested simplification as they read different parts of the article to improve understanding of challenging material. For example: *“Could you please simplify the third paragraph in the introduction?”* *After quoting the article: “Break this section of the paper down for me and explain this process in simpler terms*.*”* In general, we saw that undergraduate researchers engaged with ChatGPT in a variety of ways based on the ChatGPT-assisted reading protocol provided to them. Most students (82%, 9 of 11) followed the protocol’s recommendation of providing an introduction prior to asking questions to allow ChatGPT to provide appropriate-level responses. By far the most common type of question asked was to provide a definition, with all students asking definition questions and a total of 31 definition questions being asked from all participants. This implies that a first barrier for undergraduate researchers reading literature is encountering unfamiliar words and concepts. As discussed in Round and Campbell’s work, students reading scientific articles are likely to skim or skip unfamiliar or challenging terms, which prevents them from fully engaging with the literature.^2^ Because we observed that fewer students asked the chatbot to summarize or simplify the text, it may be worthwhile to include explicit instructions and suggestions of different ways to interact with the chatbot in future versions of the protocol. This will be explored in future studies. By providing the students with a ChatGPT-assisted reading protocol, undergraduate researchers were given a useful way to obtain definitions for unfamiliar terms and to ask follow-up questions using their natural language. The undergraduate researchers could also ask ChatGPT to simplify language when needed, further lowering the barrier to engaging with the text.

### Post-implementation survey

After completing the reading activity with the ChatGPT-assisted reading protocol, students were asked to complete a post-implementation survey about their experience using the reading protocol and their experience with ChatGPT. Eleven students completed the survey. The questions and responses are provided in Supplemental File 2.

Students were asked whether they had previously used a reading protocol and whether they had used ChatGPT before (Figure 7). We found that 73% (8 of 11) of students had not used a reading protocol before participating in the study. From the survey, 91% (10 of 11) of the students had used ChatGPT prior to this study. When asked how they used ChatGPT prior to this study in a free response question, 50% (5 of 10) reported using it to understand concepts from class and/or research, and 20% (2 of 10) reported being required to use it for class assignments. Other students reported using ChatGPT for non-academic related uses. This shows most students had familiarity with ChatGPT prior to using the ChatGPT-assisted reading protocol in this study.

**Figure 7:**
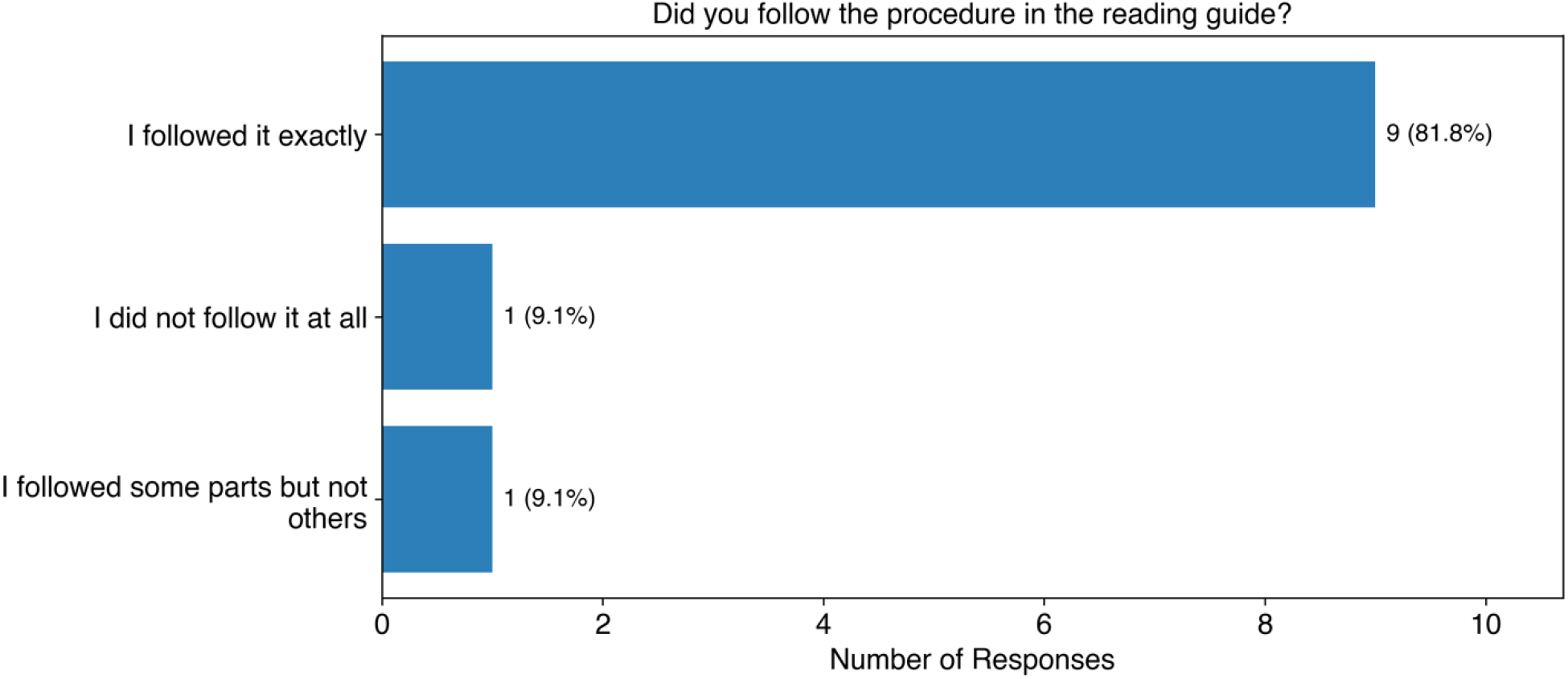
Student responses to whether they followed the reading procedure. The original question posed was “When reading the abstract and introduction to the scientific journal article with ChatGPT, did you follow the procedure in the reading guide?”

Students were also asked if they followed the procedure (Figure 7). Most students (82%, 9 of 11) followed the procedure exactly. The rest of the students either didn’t follow it at all (9%, 1 of 11), or followed some parts but not others (9%, 1 of 11). A free response was provided for students who did not follow the procedure. One student responded:

*“I did not take any notes on the side, anything I had a question about was asked to ChatGPT. “*

This is an interesting response because it suggests that the use of ChatGPT might serve as a way to interactively take notes while reading the paper, which may be worth exploring in future work.

We also asked students whether they asked clarifying or follow up questions after asking their initial question (Figure 8). None of the students said they asked follow up or clarifying questions all or most of the time. We found that 27% (3 of 11) students reported sometimes asking follow up questions, along with 27% (3 of 11) who said they asked follow up or clarification questions once or twice. We also found that 46% (5 of 11) said they never asked follow up questions, which corresponds to the results we found in the open coding of student ChatGPT transcripts. Again, this suggests to us that we should include more explicit instructions in the reading protocol so that students can consider asking different types of questions when they need to understand a concept.

**Figure 8:**
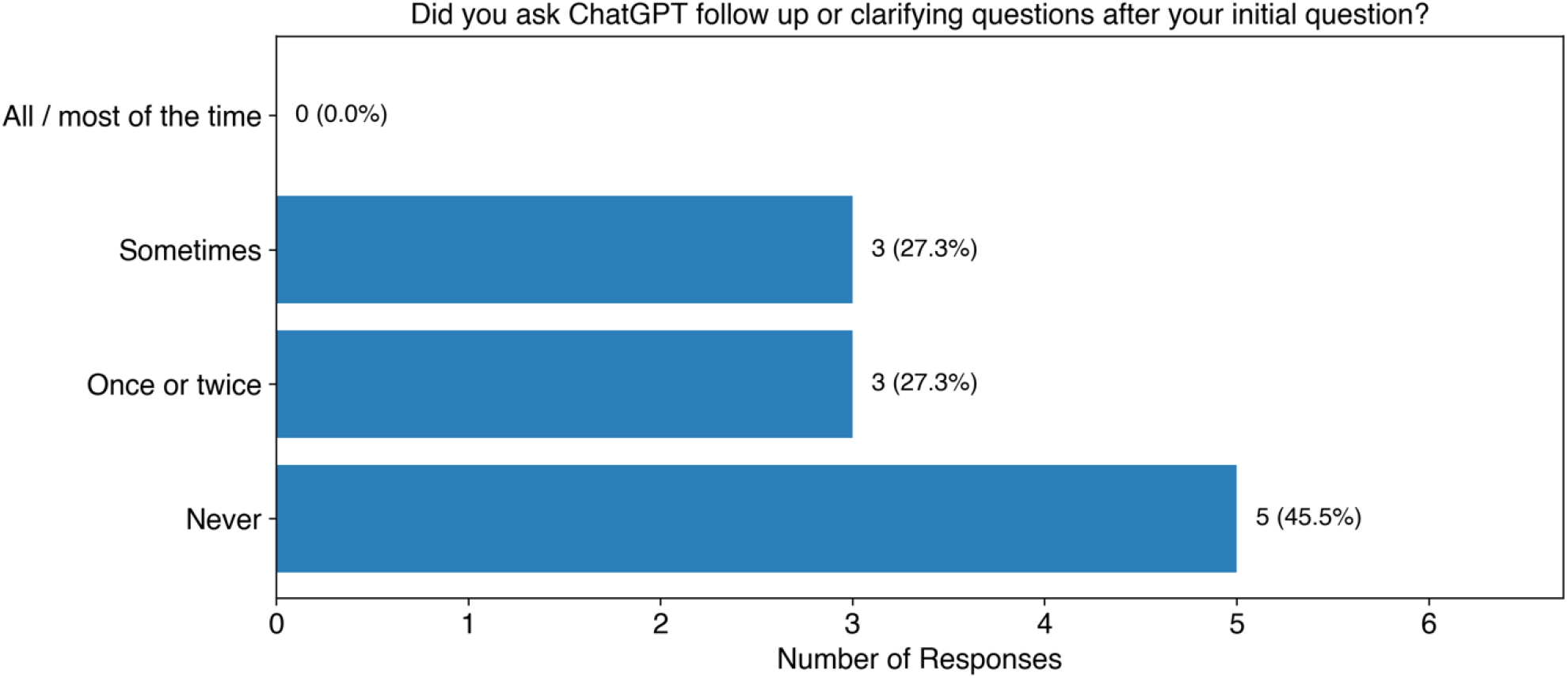
Student responses to whether they asked follow up or clarifying questions. The original question posed was “When you used ChatGPT to help with unfamiliar words, phrases, or concepts, did you ask ChatGPT follow up or clarifying questions after your initial question?”

The students were also asked to write in free response items on the biggest benefit and challenge of using ChatGPT while reading a scientific article. For the biggest benefit, the most common theme in response was that ChatGPT provides definitions and explanations in the context of the article (46%, 5 of 11), followed by clarity of explanation (36%, 4 of 11). Within their responses, students also expressed appreciation of the dynamic nature of interacting with ChatGPT (see second and third quotes) as opposed to traditional search engines:

*“Having unknown words be defined in the context of the article”*

*“The biggest benefit of using ChatGPT was not only being able to receive an answer, but also being able to tune how concise I wanted that answer to be. It’s also extremely helpful that I can ask a question in the context of a certain topic, and the computer will adjust accordingly*.*”*

*“The biggest benefit was having things broken down to me in a way that I could actually understand. I feel that when searching things up, it’s hard to find answers that actually explain without just reiterating what originally confused you*.*”*

When asked about the biggest challenge of using ChatGPT while reading scientific literature, the most common response was that it was a challenge to learn to phrase questions to get the desired response (46%, 5 of 11), and that the ChatGPT responses were sometimes too long (18%, 2 of 11):

*“The biggest challenge is figuring out how to ask ChatGPT what I want to know*.*”*

*“I feel like the biggest challenge using ChatGPT was providing enough background so that my question could be answered fully, coupled with the fact that I had to guess if I was providing enough context for a prompt*.*”*

Lastly, students were asked in a free response question if they would use ChatGPT to assist reading scientific journal articles in the future. All respondents (100%, 11 of 11) stated that they would use ChatGPT to assist them in reading articles going forward. Common themes in responses were that ChatGPT provides context/explains within the context of the article (46%, 5 of 11), and clarity of explanations and summaries (36%, 4 of 11):

*“I’d would* [sic] *expect to use ChatGPT in the future because it takes plenty of the guesswork out of contextualizing certain procedures and terms when reading papers on a research topic I am not familiar with*.*”*

*“I do expect to use ChatGPT in the future when reading scientific journal articles because it really does make the reading process much more easier, efficient, and effective. The program provides detailed, relevant, and real time explanations on questions and curiosities. It is also willing to word things differently and break down the information as much as necessary until you grasp the material. You can also provide it with your exact assumptions and thoughts on questions and material and it will respond to you and explain why you are right, wrong, or what it* [sic] *missing*.*”*

## Conclusions

Based on the feedback from undergraduate researcher students in this study, the ChatGPT-assisted reading protocol was seen as beneficial. While subjects predominantly asked ChatGPT definition-type questions while reading the introduction of the article during the study, their feedback focused on the fact that, unlike traditional search engines, ChatGPT provides definitions and explanations with respect to the context of the article. The students also valued the clarity and tunability of explanations provided by ChatGPT. The largest barrier to using ChatGPT was perceived to be formatting questions to elicit the desired response.

The finding that all the students surveyed said they would use ChatGPT in future paper reading was especially interesting. The use of a chatbot while reading unfamiliar material may be a positive intervention to help students continue to engage with biophysics literature without becoming discouraged. If a tool like ChatGPT can help students quickly find the information they need to continue reading a paper and continue asking questions until they are satisfied with their understanding, they may be encouraged to continue reading papers on a regular basis as they continue to develop expertise. Especially compared to using a search engine to find material, which can require a level of expertise to correctly frame the question and find the answer, the interactive, back-and-forth nature of the chatbot may make it easier for students to minimize the disruptions to their reading and not feel overwhelmed when reading biophysics papers. Additionally, this type of approach may be even useful in other contexts, such as helping students read textbooks or other material in their course work. In future investigations on using a ChatGPT-assisted reading protocol, additional training in effective question construction as well as different types of questions that can be asked would be beneficial to further remove barriers for undergraduate student researchers to effectively engage with literature. We also will be exploring how student interactions with the chatbot change when they read an entire journal article, especially how they use the chatbot to understand quantitative data and figures.

## Supplemental Information

The ChatGPT-assisted reading protocol is in the supplemental information.

## IRB

This study was granted exempt status by the St. John’s University IRB on 11/30/2023.

## Author Contributions

M. Sambar and G. R. Vázquez helped prepare the manuscript. A. V. Vázquez and F. X. Vázquez both designed the protocol, testing methodology, and prepared the manuscript. A. V. Vázquez designed the survey questions and performed the coding.

## Acknowledgments

Resources for this work were provided in part by the National Institutes of Health under award number NIH SC2GM131992 and NSF award CHE-2018427. All of the authors declare no competing interests.

## REFERENCES

(1) Pekrun, R.; Goetz, T.; Titz, W.; Perry, R. P. Academic Emotions in Students’ Self-Regulated Learning and Achievement: A Program of Qualitative and Quantitative Research. Educational Psychologist 2002, 37 (2), 91–105. DOI: 10.1207/S15326985EP3702_4.

(2) Round, J. E.; Campbell, A. M. Figure Facts: Encouraging Undergraduates to Take a Data-Centered Approach to Reading Primary Literature. CBE—Life Sciences Education 2013, 12 (1), 39–46. DOI: 10.1187/cbe.11-07-0057.

(3) Forest, K.; Rayne, S. Incorporating Primary Literature Summary Projects into a First-Year Chemistry Curriculum. Journal of Chemical Education 2009, 86 (5), 592. DOI: 10.1021/ed086p592.

(4) Dai, D. W.; Suzuki, S.; Chen, G. Generative AI for professional communication training in intercultural contexts: where are we now and where are we heading? Applied Linguistics Review 2024. DOI: doi:10.1515/applirev-2024-0184 (acccessed 2024-08-15).

(5) Castillo-Martínez, I. M.; Ramírez-Montoya, M. S. Research Competencies to Develop Academic Reading and Writing: A Systematic Literature Review. Frontiers in Education 2021, 5, Systematic Review. DOI: 10.3389/feduc.2020.576961.

(6) Anne Gold, R. A., Karen McNeal. Undergraduates’ Graph Interpretation and Scientific Paper Reading Shift from Novice-to Expert-like as a Result of Participation in a Summer Research Experience: A Case Study. Scholarship and Practice of Undergraduate Research 2021, 5, 7–19. DOI: 10.18833/spur/5/2/2.

(7) Lennox, R.; Hepburn, K.; Leaman, E.; van Houten, N. ‘I’m probably just gonna skim’: an assessment of undergraduate students’ primary scientific literature reading approaches. International Journal of Science Education 2020, 42 (9), 1409–1429. DOI: 10.1080/09500693.2020.1765044.

(8) Mitra, S.; Wagner, E. Introducing Undergraduates to Primary Research Literature. Journal of Chemical Education 2021, 98 (7), 2262–2271. DOI: 10.1021/acs.jchemed.0c01439.

(9) Otto, J. L.; McDowell, G. S.; Balgopal, M. M.; Lijek, R. S. Preprint Peer Review Enhances Undergraduate Biology Students’ Disciplinary Literacy and Sense of Belonging in STEM. Journal of Microbiology & Biology Education 2023, 24 (2), e00053–00023. DOI: doi:10.1128/jmbe.00053-23.

(10) Alasadi, E. A.; Baiz, C. R. Generative AI in Education and Research: Opportunities, Concerns, and Solutions. Journal of Chemical Education 2023, 100 (8), 2965–2971. DOI: 10.1021/acs.jchemed.3c00323.

(11) Hocky, G. M. Connecting molecular properties with plain language. Nature Machine Intelligence 2024, 6 (3), 249–250. DOI: 10.1038/s42256-024-00812-y.

(12) Hocky, G. M.; White, A. D. Natural language processing models that automate programming will transform chemistry research and teaching. Digital Discovery 2022, 1 (2), 79–83, 10.1039/D1DD00009H. DOI: 10.1039/D1DD00009H.

(13) White, A. D. The future of chemistry is language. Nature Reviews Chemistry 2023, 7 (7), 457–458. DOI: 10.1038/s41570-023-00502-0.

(14) Alasadi, E. A.; Baiz, C. R. Multimodal Generative Artificial Intelligence Tackles Visual Problems in Chemistry. Journal of Chemical Education 2024, 101 (7), 2716–2729. DOI: 10.1021/acs.jchemed.4c00138.

(15) Guo, Y.; Lee, D. Leveraging ChatGPT for Enhancing Critical Thinking Skills. Journal of Chemical Education 2023, 100 (12), 4876–4883. DOI: 10.1021/acs.jchemed.3c00505.

(16) Tassoti, S. Assessment of Students Use of Generative Artificial Intelligence: Prompting Strategies and Prompt Engineering in Chemistry Education. Journal of Chemical Education 2024, 101 (6), 2475–2482. DOI: 10.1021/acs.jchemed.4c00212.

(17) Yik, B. J.; Dood, A. J. ChatGPT Convincingly Explains Organic Chemistry Reaction Mechanisms Slightly Inaccurately with High Levels of Explanation Sophistication. Journal of Chemical Education 2024, 101 (5), 1836–1846. DOI: 10.1021/acs.jchemed.4c00235.

(18) Young, J. D.; Dawood, L.; Lewis, S. E. Chemistry Students’ Artificial Intelligence Literacy through their Critical Reflections of Chatbot Responses. Journal of Chemical Education 2024, 101 (6), 2466–2474. DOI: 10.1021/acs.jchemed.4c00154.

(19) Exintaris, B.; Karunaratne, N.; Yuriev, E. Metacognition and Critical Thinking: Using ChatGPT-Generated Responses as Prompts for Critique in a Problem-Solving Workshop (SMARTCHEMPer). Journal of Chemical Education 2023, 100 (8), 2972–2980. DOI: 10.1021/acs.jchemed.3c00481.

(20) Pence, H. E.; Hightower, G.; Forlenza, J.; Leonard, K.; McLellan, A.; Suero, A.; Amoah, B.; Mbow, M.; Borner, S.; Castillo, A.; et al. Using Generative AI Systems for Critical Thinking Engagement in an Advanced Chemistry Course: A Case Study. Journal of Chemical Education 2024. DOI: 10.1021/acs.jchemed.4c00242.

(21) McGuire, S. Y. Teach Students How to Learn: Strategies You Can Incorporate Into Any Course to Improve Student Metacognition, Study Skills, and Motivation; Routledge, 2015. DOI: 10.4324/9781003447313.

(22) Kirchdoerfer, R. N.; Wang, N.; Pallesen, J.; Wrapp, D.; Turner, H. L.; Cottrell, C. A.; Corbett, K. S.; Graham, B. S.; McLellan, J. S.; Ward, A. B. Stabilized coronavirus spikes are resistant to conformational changes induced by receptor recognition or proteolysis. Scientific Reports 2018, 8 (1), 15701. DOI: 10.1038/s41598-018-34171-7.

(23) Saldaña, J. The Coding Manual for Qualitative Researchers; SAGE Publications Ltd, 2021.

