## Supplemental Information for "A ChatGPT Assisted Reading Protocol for Undergraduate Research Students"

As part of the study, participants were provided with a ChatGPT-assisted literature reading protocol, shown in Figure S1. The goal of this protocol was to give the students a structured approach to reading the provided paper and give them specific guidance on how to incorporate ChatGPT queries into their reading. The reading protocol also included a short explanation at the end explaining to students to reasons for providing them with a structured reading approach.

Students were given an initial survey to determine their experience and confidence when reading scientific papers. An anonymized version of the questions and the responses are provided in Supplemental File 1. Students were also given a post-study survey. The questions and responses are provided in Supplemental File 2.

### Guidelines for Reading Scientific Papers

#### Previewing the Paper

1. **Understand the Title**: Determine the main focus of the paper based on its title.
2. **Analyze the Abstract**: Read the abstract, and if available, the accompanying figure, to identify the paper's major findings.
3. **Abstract Structure**: While reading the abstract, consider the typical structure: Importance, Problem, Approach, and Insight.
4. **Review Headings and Figures**: Skim through the paper's headings and figures to anticipate the questions the paper aims to answer.

#### Reading the Paper

1. **Highlight Key Text**: Use an erasable method (e.g., pencil underlining) to highlight text that seems important or intriguing. You can refine this later.
2. **Margin Notes**: Whenever you have a question, insight, or idea, jot it down in the margins. These notes are for your personal use.
3. **Extended Notes**: If you need more space for notes or sketches, use a separate notebook or sheet of paper. The goal is to capture all your thoughts for later reference.

#### Utilizing ChatGPT for Clarifications

4. **When you encounter an unfamiliar concept, term, or methodology that feels like a stumbling block, use ChatGPT to help clarify the concept**:
   a. **Start a New Conversation**: Initiate a new conversation with ChatGPT. Use this same thread for all questions related to this paper.
   b. **Introduce Yourself**: Briefly describe your academic background, lab experience, and comfort level with the material and reading scientific papers.
   c. **Inform ChatGPT About the Paper**: Provide the paper's title, field, and any other relevant information to help ChatGPT understand the context.
   d. **Ask Your Questions**: Pose your question to ChatGPT. If the initial response is unclear, continue asking follow-up questions until you get the information you're seeking.
   e. **Seek Further Clarification**: If something remains unclear, ask for additional context or a different type of explanation.
   f. **Explore Further**: If a point seems particularly relevant or interesting, feel free to continue the conversation with ChatGPT as long as needed to fully understand the paper.
5. **Post-Reading Summary**: After completing the paper, review your notes and write a brief summary that you can refer to later. This can be as short as one sentence.

#### Objective

The primary aim of this method is to facilitate a smooth reading experience, minimizing disruptions to your flow. The preview stage primes you for the paper's content, while the reading stage aims to maintain a steady pace. The use of ChatGPT serves as a real-time resource for overcoming stumbling blocks in comprehension.

**Figure S1**: ChatGPT-assisted literature reading protocol provided to study participants.
